# Mitogenome Diversity and Phylogenetic Insights of *Aedes albopictus* in Greece

**DOI:** 10.1101/2024.12.20.629447

**Authors:** Georgios Balatsos, Venetia Karathanasi, Vasiliki Evangelou, Despoina Κapadaidaki, Nikolaos Tegos, Anastasia Panagopoulou, Marina Bisia, Vasileios Karras, Dimitrios P. Papachristos, Nikos T. Papadopoulos, Antonios Augustinos, Elina Patsoula, Antonios Michaelakis

## Abstract

The invasive mosquito species *Aedes albopictus*, commonly known as the Asian Tiger Mosquito, is a significant global health concern due to its role as a vector for diseases such as Dengue, Zika, and Chikungunya. In Greece, Ae. albopictus was first detected in 2005, with subsequent widespread establishment across the country. This study investigates the genetic diversity and phylogeography of Greek Ae. albopictus populations by analyzing mitochondrial cytochrome oxidase I *(COI)* gene sequences. A total of 488 female individuals were analyzed, revealing six haplotypes in samples collected from 2016 to 2018 using the LCO 1490 and HCO 2198 primer pair, including instances of identical haplotypes across years, while only one haplotype was consistently detected across all 2019 to 2022 samples using the UBC6 and UBC9 primer pair. These findings underscore the importance of molecular tools in understanding invasion dynamics and informing targeted surveillance and control measures. Further research is needed to assess additional worldwide populations and expand phylogeographic comparisons to elucidate global dispersal patterns.

## Introduction

*Aedes (Ae*.*) albopictus*, also known as the “Asian Tiger Mosquito”, is listed among the 100 most invasive species globally. It is a highly efficient vector of several significant human diseases, including Dengue [1,2], Zika, Chikungunya virus [3], Yellow fever [4], Japanese encephalitis [5,6] and zoonoses, such as dirofilarioses [7,8] *Ae. albopictus* native region are the tropical forests of Southeast Asia, but the last 34 years through human travel and commerce it has spread to Africa, Europe, Australia and America [2,9], namely to every continent except Antarctica. The widespread distribution of *Ae. albopictus* outside its native home - range is presumed to have been primarily human-mediated and accidental [10]. Due to its ability to colonize a wide range of natural and artificial breeding places together with the resistance of its eggs to desiccation and its relative lack of host specificity [11], the species has successfully competed with co - occurring mosquito species and has been able to rapidly build up large populations in new geographic regions and has adapted to a variety of different environmental conditions [12–14].

In Europe, after its first detection in Albania in 1979 [15,16], the species was found in Genoa in September 1990 [17]. This latter introduction is considered to have resulted from tire imports from the United States [17–19]. Since then, *Ae. albopictus* has spread throughout the entire mainland of Italy as well as Sicily and Sardinia [18–20]. In France, *Ae. albopictus* was first reported in Normandy in 1999 [21]. Nowadays, established homogenous populations of the species occur in all countries of the Mediterranean Sea, including parts of Turkey and the Middle Eastern states of Lebanon, Israel and Syria [22,23]. Italy and southern France are the most infested regions, since *Ae*.*albopictus* has been established in most areas of these countries [23]. Moreover, *Ae. albopictus* populations have been newly introduced into regions in Cyprus, Czechia, Liechtenstein, the Netherlands, Slovakia, Slovenia, Spain, Portugal and Sweden [24,25].

In Greece, the presence of *Ae. albopictus* was first reported in 2005 in the northwest by Samanidou-Voyadioglou et. al.2005 [26]. By 2006, the species had established populations in various parts of the country, including the island of Corfu and mainland Thesprotia. Following its initial detection, *Ae. albopictus* exhibited slow dispersal patterns until 2010, after which it underwent a rapid and widespread expansion, reaching regions such as Central Macedonia, Peloponnese, and Attica. By 2024, *Ae. albopictus* had been detected throughout most of regions of the country, with the exception of some areas in northern Greece (Region of West Macedonia) and certain Aegean islands where data are lacking [27–30].

*Aedes albopictus* is not only a nuisance species with significant impacts on environmental health and community welfare but also a highly competent vector of numerous arboviruses and parasites, posing serious concerns for both veterinary and public health. In the absence of *Ae. aegypti* in European, *Ae. albopictus* has become the primary vector of significant arboviruses in the region. Its role in the transmission and spread of pathogenic flaviviruses (e.g., Dengue, Zika, West Nile, Yellow fever, and Japanese encephalitis), alphaviruses (e.g., Chikungunya), and bunyaviruses (e.g., La Crosse and Rift Valley fever) underscores its importance as a major global public health threat. The emergence of indigenous vector-borne diseases in recent years, such as Chikungunya and Dengue, is closely associated with the distribution and activity of *Ae. albopictus*. Chikungunya virus outbreaks in Europe have been sporadically reported since 2007, when a major outbreak occurred in the Emilia Romagna region of Italy, resulting in 330 autochthonous cases [31]. Subsequent cases were recorded in France during 2010, 2014, and 2017, and in Italy again in 2017, with 270 cases being reported [32–38]. No additional Chikungunya cases have been documented in these countries up to 2024. Autochthonous Dengue cases, caused by serotypes 1 and 2, have also been documented in Europe over the past 14 years, primarily in Croatia, France, Spain, and Italy. In Croatia, ten cases were reported in 2010, with no further cases recorded thereafter. France reported its first two autochthonous cases in 2010, followed by sporadic cases in 2013, 2014, 2015, 2018, 2019, 2020, and 2021 [33,39,40]. However, a notable increase was observed from 2022 onwards, with 65 cases reported in 2022, 45 in 2023, and 83 in 2024 [41,42]. Spain documented its first six autochthonous cases in 2022, followed by three cases in 2023 and eight in 2024 [42,43]. In Italy, 10 cases were recorded in 2020, followed by a significant outbreak with 82 cases in 2023 and 213 cases in 2024 [43–45].

Since 2014, an extensive oviposition network has been established in Greece to study invasive mosquito species and this effort was conducted within the framework of projects during the years. The network was developed following the protocol described by Giatropoulos et al. 2012 [27] and was strategically focused on specific regions to implement the oviposition network and conduct molecular analyses. These regions were selected as potential points of entry for the species into Greece, given their air or land connections with neighboring countries where the species had already been detected.

Specific molecular markers are essential for monitoring the spread, local adaptation, and evolution of *Ae. albopictus* populations, as well as for gaining deeper insights into their population structure. Details of population genetics and structure will subsequently allow and possibly predict the geographical and temporal dynamics of the species expansion. This is a fundamental requirement both for the development of strict monitoring protocols and for the improvement of sustainable control measures and practical operations of control programs.

The investigation of variation in mitochondrial and nuclear DNA offers an effective tool for assessing the phylogeographic history of an organism, especially when samples are available not only from the area of introduction but also from the area of origin. Previous studies have already examined phylogeographic relationships among different populations of *Ae. albopictus* from all over the world, using mitochondrial and nuclear markers [46–50] and provided evidence that in this species, mitochondrial genes display mediate to high levels of genetic variation and sequence divergence. Among them, there is only one small – scale DNA - based study conducted up to date investigating the genetic diversity of Greek *Ae*.*albopictus* populations, however it is limited to only a few COI sequences from two geographic regions of Greece[51]. The population genetics of this species, however, merit further exploration and additional sampling is a prerequisite in order to confirm the already identified patterns and to discover new possible ones. The objectives of this study were: (1) to determine the pattern of genetic variability within the Greek *Ae. albopictus* populations based on analyses of the sequence diversity of the mitochondrial gene cytochrome oxidase I (COI) and (2) to establish the evolutionary relationships of these Greek populations with other *Ae. albopictus* worldwide populations and (3) to determine the geographic origin of populations of the species that colonized Greece.

## Materials and methods

### Mosquito samples

Ovitraps were placed at sampling sites across the country and the duration of sampling was weekly. The wooden substrates were collected and transported to BPI, ensuring strict adherence to the protocol guidelines for the operation and maintenance of the ovitraps. In each one of the aforementioned regions, one to nine distinct locations were selected for placing ovitraps. The individuals which were collected from 2014 until 2022 during the entomological surveillance and emerged from the ovitraps were singly used for molecular analysis. All specimens included in this study, their code number, the date of collection and the location of collection site are shown in detail in Table 1. The wooden substrates from ovitraps were collected eggs that were laid attached to water-filled ovitraps. Egg hatching was carried out in 1-liter beakers containing 700 ml of water and 2 ml of a deionized water solution consisting of 12.5% w/v nutrient broth powder (Oxoid, UK) and 2.5% w/v yeast extract powder (Oxoid, UK) [52]. Rearing was performed under controlled temperature (25±2 °C), relative humidity (65±5%), and lighting conditions (L14:D10, with sunset and sunrise simulation for 30 mins, respectively). First instar larvae were placed in large plastic containers supplied with a daily provision of fish food (0.1 mg/larva) (“NovoTom Artemia”, JBL, Germany) until the pupal stage. Developed pupae were collected daily and transferred to another plastic containers where they developed into adult mosquitoes. Emerged adult mosquitoes were identified by using published literature [53,54] and used for the subsequent analyses.

**Table 1.** Number of specimens collected across different regions in Greece from 2014 to 2022.

| Region | Area | No. specimens per year |  |  |  |  |  |  |  |  |
| --- | --- | --- | --- | --- | --- | --- | --- | --- | --- | --- |
|  |  | 2014 | 2015 | 2016 | 2017 | 2018 | 2019 | 2020 | 2021 | 2022 |
| ATTICA | KIFISIA |  |  |  |  |  |  |  | 20 | 20 |
| ATTICA | ATHENS AIRPORT | 25 | 25 | 13 | 21 | 20 | 20 | 16 |  |  |
| **ATTICA** | ATHENS PORT | 25 | 25 | 5 | 17 | 20 | 20 | 16 |  |  |
| **ATTICA** | RIZOUPOLI | 25 |  |  |  |  |  |  |  |  |
| **CRETE** | CHANIA (airport & port) |  | 25 | 21 | 20 | 20 | 20 | 16 | 20 | 20 |
| **CRETE** | RETHYMNO |  |  | 23 | 22 | 20 | 20 |  |  |  |
| **CRETE** | HERAKLEION |  | 12 |  |  |  |  |  |  |  |
| **East Macedonia and Thrace** | KAVALA |  |  | 10 | 20 | 20 |  | 16 | 20 | 20 |
| **North Aegean** | MYTILENE |  |  | 6 | 3 | 20 |  |  |  |  |
| **Central Macedonia** | THESSALONIKI (airport & port) |  | 1 | 1 |  | 29 | 20 |  |  |  |
| **East Macedonia and Thrace** | DRAMA |  |  |  | 2 |  |  |  |  |  |
| **South Aegean** | SYROS |  |  |  |  |  | 3 |  |  |  |
| **Thessaly** | LARISA |  |  | 0 | 0 | 0 | 0 | 0 | 20 | 20 |
| **Total number of specimens** |  | 75 | 88 | 79 | 105 | 149 | 103 | 64 | 80 | 80 |

**Figure 1.**
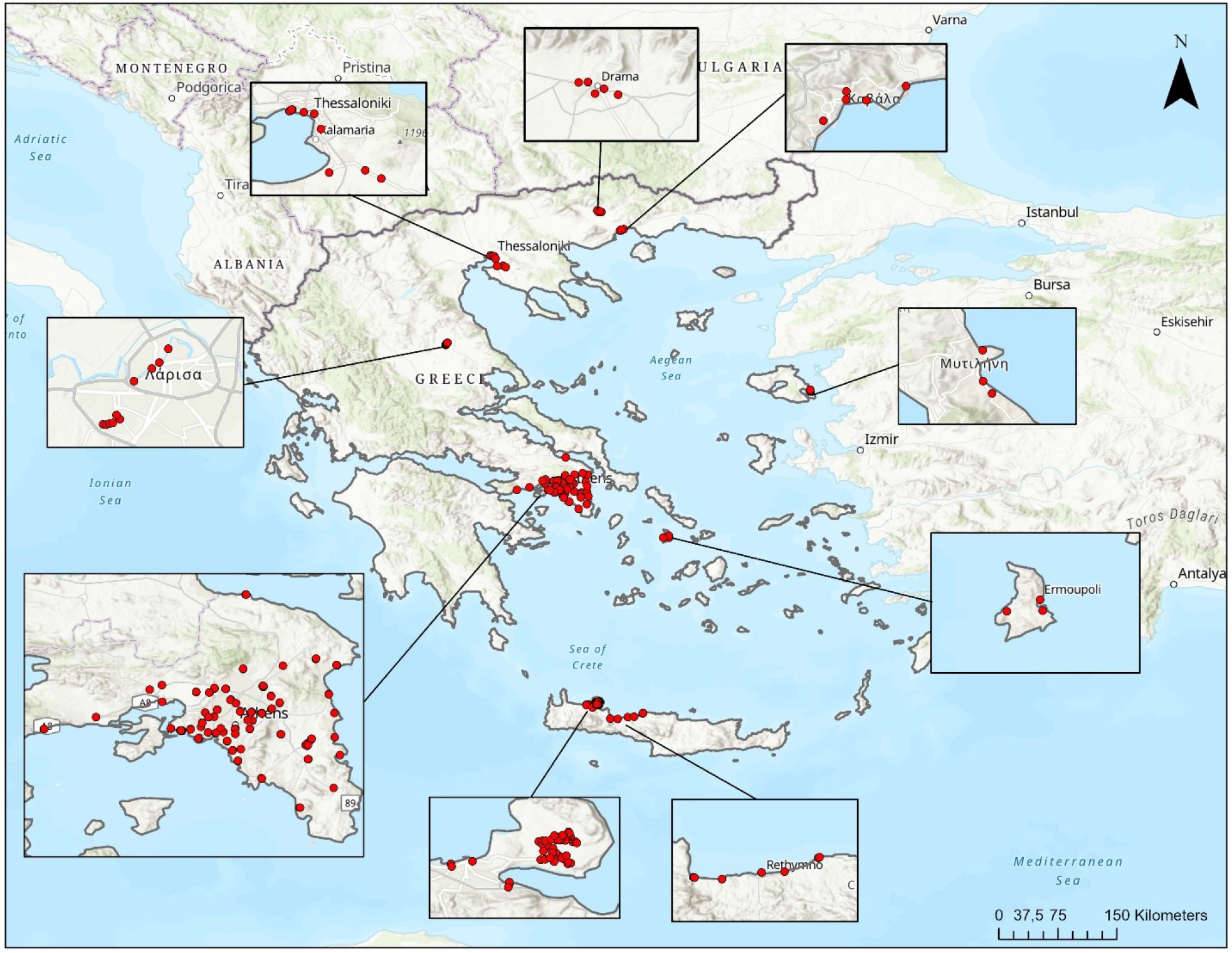
Oviposition network in Greece during the years, 2014 to 2022.

### DNA extraction and PCR

For the molecular analysis of the Greek mosquito populations, the emerged female adults were collected and were immediately preserved in 98% ethanol at -20°C until further processing. Total genomic DNA (gDNA) was extracted from the whole body of single adult individuals by applying different extraction protocols during the years. Partial sequences of the mitochondrial DNA (mtDNA) gene encoding for the cytochrome oxidase I *(COI)* gene were obtained through Polymerase Chain Reaction (PCR).

### 2014 – 2015

In 2014, *Ae. albopictus* specimens were collected from multiple locations in Greece, including the Athens port, Athens airport, Rizoupoli (the area of the first detection of *Ae. albopictus* in Athens), and the Thessaloniki port. In 2015, *Ae. albopictus* was recorded for the first time on the island of Crete [55] and therefore specimens were collected from Chania and Heraklion. Additional specimens were also collected that year from the Athens airport and Athens port. Genomic DNA was extracted using the PureLine® Genomic DNA Kit (Invitrogen, Waltham, MA, USA), following the manufacturer’s protocol. Polymerase chain reaction (PCR) amplification was run in a final volume of 25 μl using the primers JERRY : 5’-CAA CATTTATTT TGA TTT TTT GG-3’ and TL2-N-3014 PAT: 5’-TCC AAT GCACTA ATC TGC CAT ATT 3’ [56] that amplify a 825bp fragment of the mitochondrial *COI* gene. Each reaction contained 8 μl of the extracted DNA, 10.6 μl of double distilled water, 5 μl of Red Bioline buffer (provided with the Taq), 0.5 μl of primer Jerry, 0.5 μl of primer Pat, and 0.4 ll of MyTaq (Red Bioline). PCR procedure was as follows: 3 min of 94 °C, followed by 40 cycles of 30 s at 94 °C (denaturation), 30 s at 45 °C (annealing), and 1.5 min at 72 °C (extension). The final extension period was carried out at 72 °C for 7 min.

### 2016-2018

For the individuals collected from 2016 to 2018, DNA extraction was implemented with different protocols to test their efficiency and the different yields of DNA extracts they produce. The commercially available kits, NucleoSpin DNA insect and NucleoSpin Tissue (Macherey – Nagel, Germany) were used according to the manufacturers’ instructions. In addition, the cetyltrimethyl ammonium bromide (CTAB) DNA extraction method was tested and performed as previously described [57,58]. The *COI* gene was amplified using the primers LCO - 1490 (5’ - GGTCAACAAATCATAAAGATATTGG - 3’) and HCO – 2198 (5’ – TAAACTTCAGGGTGACCAAAAAATCA – 3’) [59]. Two microliters of the gDNA extract were used as the template in 20μl reactions containing 0.2 mM dNTPs, 1.0 μM of each primer, 1 Kapa HiFi Taq DNA polymerase (Kapa Biosystems, Cape Town, South Africa) and 1x enzyme buffer.

PCRs were implemented under the following conditions: one step of initial denaturation at 95 °C for 3 min followed by 40 cycles at 95 °C for 30 sec, 48 °C for 30 sec and 72 °C for 1 min. Final extension was performed at 72 °C for 2 min.

### 2019-2022

Specimens collected from 2019 until 2022, DNA extraction was conducted using Maxwell 16 Automated Nucleic Acid extraction system (Promega, Madison, WI, USA) with the Maxwell 16 LEV. PCRs were conducted using the primers UBC6 (5’-GGA GGA TTT GGA AAT TGA TTAGTT CC-3’) - UBC9 (5’-CCC GGT AAA ATTA AAA ATA TAA ACT TC-3’) [56] which amplify a 474 bp fragment of the *COI* gene. Two microliters of the gDNA extract were used as the template in 20μl reactions containing 0.2 mM dNTPs, 1.0 μM of each primer, 1 Kapa HiFi Taq DNA polymerase (Kapa Biosystems, Cape Town, South Africa) and 1x enzyme buffer. PCRs were implemented under the following conditions: one step of initial denaturation at 94 °C for 5 min followed by 40 cycles at 94 °C for 60 sec, 50 °C for 60 sec and 72 °C for 60 sec. Final extension was performed at 72 °C for 5 min.

### Sequencing

All obtained pcr products from 2016 to 2022 were visualized on a 1.2% agarose gel electrophoresis containing Midori Dye, Green Staining and were purified using the NucleoFast PCR Clean-up kit (Macherey – Nagel, Germany) according to the manufacturer’s instructions. Purification of PCR products was performed in both directions using the primers mentioned above by Macrogen sequencing service (Amsterdam, The Netherlands). and CEMIA SA. (Larissa, Greece). Sequences obtained in the present study were analyzed using Geneious Prime software (https://www.geneious.com/) and were compared with the corresponding ones available at GenBank using the BLAST algorithm of National Center for Biotechnology Information (NCBI, http://www.ncbi.nlm.nih.gov).

## Results

In total, 488 female individuals from 823 were analyzed for examining the genetic diversity of the mtCOI gene. Among the 57, 37 and 98 individuals which were collected in 2016, 2017 and 2018, respectively, and were analyzed with the primer pair LCO-1490 / HCO-2198, 6 haplotypes were detected (one in 2016, one in 2017 and four in 2018) (Hap_Aea_2016-1, Hap_Aea_2017-1 Hap_Aea_2018-1, Hap_Aea_2018-2, Hap_Aea_2018-3, Hap_Aea_2018-4). Haplotypes Hap_Aea_2016-1 and Hap_Aea_2018-4 were identical and haplotypes Hap_Aea_2017-1 and Hap_Aea_2018-1 were identical, also (100% identity in the overlapping fragment of 658 bp). Among the 100 individuals collected in 2019, the 38 in 2020, the 78 in 2021 and the 80 in 2022, that were tested using the primer pair UBC6/UBC9, one haplotype was identified (Hap_Aea_2019-1, Hap_Aea_2020-1, Hap_Aea_2021-1, Hap_Aea_2022-1) (100% identity in the overlapping fragment of 472 bp).

**Figure 2.**
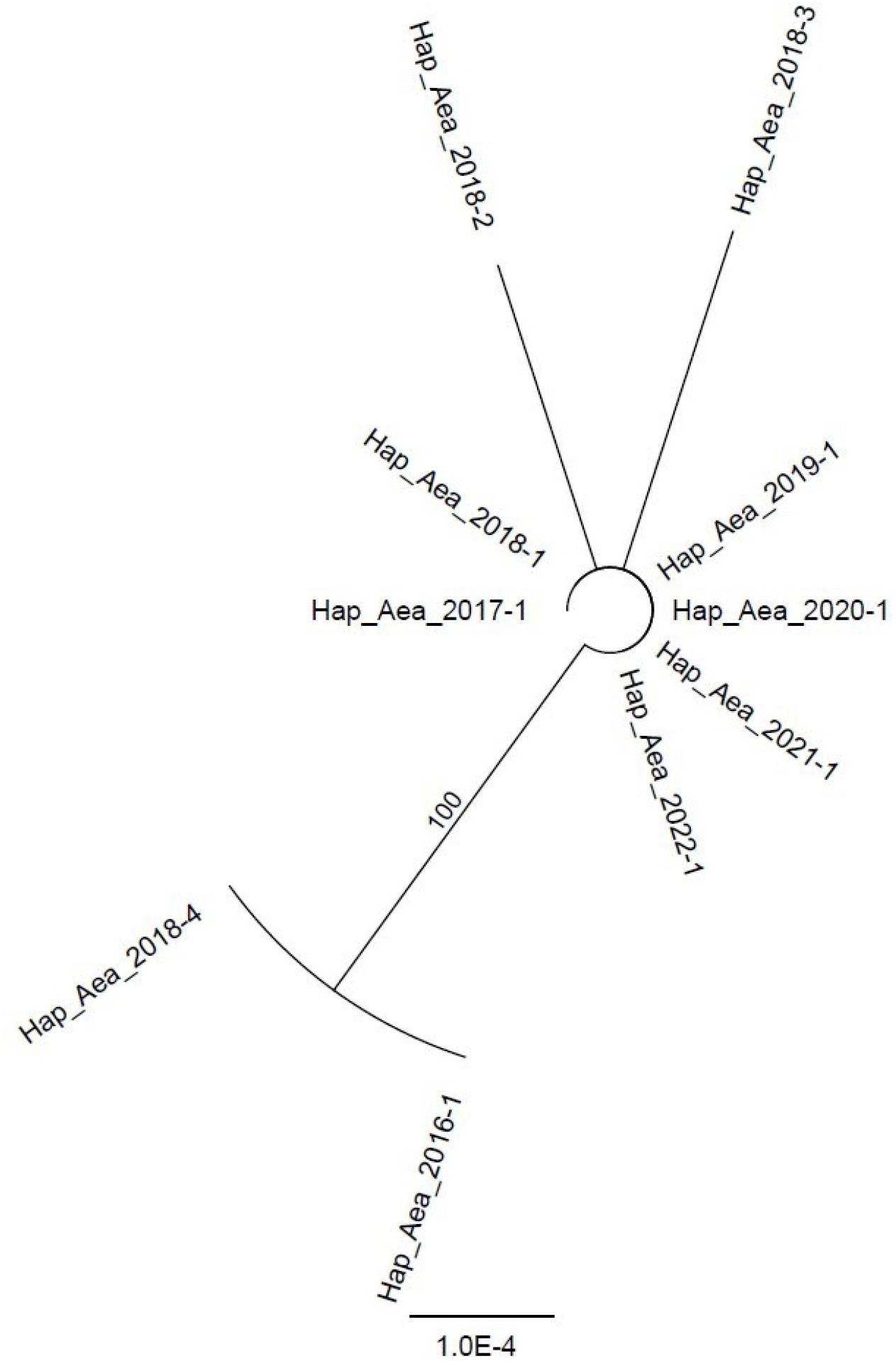
Phylogenetic reconstruction of Greek Aedes albopictus populations obtained from the analysis of *COI* sequences. Numbers above lines indicate posterior probabilities (%) of Bayesian Inference (only when above 50%)..

## Discussion

In this study we have exploited the mitochondrial marker *COI* (cytochrome oxidase I) that has allowed us to highlight the presence of the intraspecific variability and the level of differentiation of *Ae. albopictus* populations collected in different areas of Greece. The results presented here, gave an insight of the distribution of genetic diversity within and between populations and showed that most of the genetic variation was detected at the individual level.

According to our results, we suggest that the genetic diversity identified in our study is a consequence of several independent introduction events of the species, each one corresponding to one different haplotype, followed by their expansion throughout the whole country. The haplotype variation, has a distribution that is independent from geography. This genetic feature and differentiation pattern has been also revealed by using other markers and by examining populations of the species from other countries [60–64].

The mitochondrial marker here implemented, represent an important tool and provide several clues regarding untangling the possible routes of invasion, the identification of the origins of mosquito populations and the dispersion dynamics of this species. The mitochondrial cytochrome oxidase subunit 1 *(COI)* gene is one of the most popular markers used for molecular systematics in mosquitoes. It can be used to track divergence in very closely related taxa and even within species. Fragments of the gene are used to infer phylogenies and a significant amount of sequences are deposited in the DNA barcoding database. Therefore it is important to understand the evolutionary relationship of the *COI* gene among and within mosquito species. Due to the fact that mitochondrial genes are present in multiple copies and have the advantage of being maternally inherited [65], they have been widely used in studies of population genetics. Cytochrome oxidase I *(COI)* is one of the most variable genes of mitochondrial DNA and has been studied in several species of the Anopheles genera.

The threat represented by this mosquito species is growing due to the lack of sustainable control measures and of the progressive spread of insecticide – resistance [66,67]. The detailed knowledge of the population structure and of the molecular basis of the genomic flexibility of the vector species and hence of the mechanisms ensuring the diversity of its populations [68,69] is crucial and can be used in future efficient surveillance methods and control programs. Therefore, it appears necessary to examine more individuals from more regions across the country in order to assure the existence of the detected haplotypes and determine the existence of new ones. In addition, the comparison with other sequences should be continued in order to have a complete picture of the phylogeography and distribution of Ae. albopictus’ populations.

## Acknowledgments

We extend our gratitude to Evangelia Zavitsanou, Geographer at the Scientific Directorate of Entomology and Agricultural Zoology, Benaki Phytopathological Institute, for her valuable contribution in producing the map featured in this manuscript.

## Authors’ contributions

Conceptualization GB, AM, EP; Data curation, GB, DK, VE, AP, NT, MB, VK; Formal analysis, DK,VE, EP, NT, VK Funding acquisition, AM.;EP Methodology, GB, DK, VK, AP, VE, VKar, NT, MB, DP; Project administration, AM, EP; Resources, AM, EP; Supervision, NP, AM, EP; Writing – original draft, GB, AM, EP; DK Writing – review & editing, GB, DK, VK, VE, NT, VKar, MB, NP, AM, EP. All the authors have read and agreed to the published version of the manuscript.

## References

1. Lundström, J.O. Mosquito-Borne Viruses in Western Europe: A Review. J Vector Ecol 1999, 24, 1–39.

2. Benedict, M.Q.; Levine, R.S.; Hawley, W.A.; Lounibos, L.P. Spread of the Tiger: Global Risk of Invasion by the Mosquito Aedes Albopictus. Vector Borne Zoonotic Dis 2007, 7, 76–85, doi:10.1089/vbz.2006.0562.

3. Schaffner, F.; Medlock, J.M.; Van Bortel, W. Public Health Significance of Invasive Mosquitoes in Europe. Clin Microbiol Infect 2013, 19, 685–692, doi:10.1111/1469-0691.12189.

4. Countries with Risk of Yellow Fever Transmission and Countries Requiring Yellow Fever Vaccination (November 2022) Available online: https://www.who.int/publications/m/item/countries-with-risk-of-yellow-fever-transmission-and-countries-requiring-yellow-fever-vaccination-(november-2022) (accessed on 18 December 2024).

5. Zeller, H. Is Japanese Encephalitis Emerging in Europe? Eurosurveillance 2012, 17, 20242, doi:10.2807/ese.17.32.20242-en.

6. Gossner, C.M.; Dhollander, S.; Presser, L.D.; Briet, O.; Bakonyi, T.; Schaffner, F.; Figuerola, J. Potential for Emergence of Japanese Encephalitis in the European Union. Zoonoses and Public Health 2024, 71, 274–280, doi:10.1111/zph.13103.

7. Cancrini, G.; Frangipane di Regalbono, A.; Ricci, I.; Tessarin, C.; Gabrielli, S.; Pietrobelli, M. Aedes Albopictus Is a Natural Vector of Dirofilaria Immitis in Italy. Veterinary Parasitology 2003, 118, 195–202, doi:10.1016/j.vetpar.2003.10.011.

8. Capelli, G.; Frangipane di Regalbono;, Simonato, G. A.; Cassini, R.; Cazzin, S.; Cancrini, G.; Otranto, D.; Pietrobelli, M. Risk of Canine and Human Exposure to Dirofilaria Immitis Infected Mosquitoes in Endemic Areas of Italy. Parasites & Vectors 2013, 6, 60, doi:10.1186/1756-3305-6-60.

9. Madon, M.B.; Mulla, M.S.; Shaw, M.W.; Kluh, S.; Hazelrigg, J.E. Introduction of Aedes Albopictus (Skuse) in Southern California and Potential for Its Establishment. J Vector Ecol 2002, 27, 149– 154.

10. Tatem, A.J.; Hay, S.I.; Rogers, D.J. Global Traffic and Disease Vector Dispersal. Proc Natl Acad Sci U S A 2006, 103, 6242–6247, doi:10.1073/pnas.0508391103.

11. Hawley, W.A. The Biology of Aedes Albopictus. J Am Mosq Control Assoc Suppl 1988, 1, 1–39.

12. Lounibos, L.P. Invasions by Insect Vectors of Human Disease. Annu Rev Entomol 2002, 47, 233– 266, doi:10.1146/annurev.ento.47.091201.145206.

13. Armistead, J.S.; Arias, J.R.; Nishimura, N.; Lounibos, L.P. Interspecific Larval Competition between Aedes Albopictus and Aedes Japonicus (Diptera: Culicidae) in Northern Virginia. J Med Entomol 2008, 45, 629–637, doi:10.1603/0022-2585(2008)45[629:ilcbaa]2.0.co;2.

14. Medley, K.A. Niche Shifts during the Global Invasion of the Asian Tiger Mosquito, Aedes Albopictus Skuse (Culicidae), Revealed by Reciprocal Distribution Models. Global Ecology and Biogeography 2010, 19, 122–133, doi:10.1111/j.1466-8238.2009.00497.x.

15. Adhami, J.; Reiter, P. Introduction and Establishment of Aedes (Stegomyia) Albopictus Skuse (Diptera: Culicidae) in Albania. J Am Mosq Control Assoc 1998, 14, 340–343.

16. Adhami: Mushkonjat (Diptera: Culicidae) Te Shqiperise,… - ?ελετητής Google Available online: https://scholar.google.com/scholar_lookup?journal=Revista%20Mjek%C3%ABsore&title=Mushkonjat%20(Diptera:%20Culicidae)%20te%20Shqiperise,%20tribu%20Culicini&author=J%20Adhami&volume=1%E2%80%932&publication_year=1987&pages=82-95&#d=gs_cit&t=1734506114844&u=%2Fscholar%3Fq%3Dinfo%3ANU6vXzRBQR0J%3Ascholar.google.com%2F%26output%3Dcite%26scirp%3D0%26hl%3Del (accessed on 18 December 2024).

17. Sabatini, A.; Raineri, V.; Trovato, G.; Coluzzi, M. [Aedes albopictus in Italy and possible diffusion of the species into the Mediterranean area]. Parassitologia 1990, 32, 301–304.

18. Dalla Pozza, G.; Majori, G. First Record of Aedes Albopictus Establishment in Italy. J Am Mosq Control Assoc 1992, 8, 318–320.

19. Dalla Pozza, G.L.; Romi, R.; Severini, C. Source and Spread of Aedes Albopictus in the Veneto Region of Italy. J Am Mosq Control Assoc 1994, 10, 589–592.

20. Knudsen, A.B. Geographic Spread of Aedes Albopictus in Europe and the Concern among Public Health Authorities. Report and Recommendations of a Workshop, Held in Rome, December 1994. Eur J Epidemiol 1995, 11, 345–348, doi:10.1007/BF01719441.

21. Schaffner, F.; Karch, S. Première Observation d’Aedes Albopictus (Skuse, 1894) En France Métropolitaine. Comptes Rendus de l’Académie des Sciences - Series III - Sciences de la Vie 2000, 323, 373–375, doi:10.1016/S0764-4469(00)00143-8.

22. Caminade, C.; Medlock, J.M.; Ducheyne, E.; McIntyre, K.M.; Leach, S.; Baylis, M.; Morse, A.P. Suitability of European Climate for the Asian Tiger Mosquito Aedes Albopictus: Recent Trends and Future Scenarios. J R Soc Interface 2012, 9, 2708–2717, doi:10.1098/rsif.2012.0138.

23. Medlock, J.M.; Hansford, K.M.; Schaffner, F.; Versteirt, V.; Hendrickx, G.; Zeller, H.; Van Bortel, W. A Review of the Invasive Mosquitoes in Europe: Ecology, Public Health Risks, and Control Options. Vector Borne Zoonotic Dis 2012, 12, 435–447, doi:10.1089/vbz.2011.0814.

24. Zé-Zé, L.; Borges, V.; Osório, H.C.; Machado, J.; Gomes, J.P.; Alves, M.J. Mitogenome Diversity of Aedes (Stegomyia) Albopictus: Detection of Multiple Introduction Events in Portugal. PLOS Neglected Tropical Diseases 2020, 14, e0008657, doi:10.1371/journal.pntd.0008657.

25. ECDC 2024 European Centre for Disease Prevention and Control and European Food Safety Authority. Mosquito Maps.

26. Samanidou-Voyadjoglou, A.; Patsoula, E.; Spanakos, G.; Vakalis, N.C. European Mosquito Bulletin, 19 (2005), 10–11. Journal of the European Mosquito Control Association ISSN1460-6127.

27. Giatropoulos, A.; Emmanouel, N.; Koliopoulos, G.; Michaelakis, A. A Study on Distribution and Seasonal Abundance of Aedes Albopictus (Diptera: Culicidae) Population in Athens, Greece. Journal of Medical Entomology 2012, 49, 262–269, doi:10.1603/ME11096.

28. Beleri, S.; Chatzinikolaou, S.; Nearchou, A.; Patsoula, E. Entomological Study of the Mosquito Fauna in the Regional Unit of Drama, Region of East Macedonia-Thrace, Greece (2015 to 2016). Vector-Borne and Zoonotic Diseases 2017, 17, 665–671, doi:10.1089/vbz.2017.2113.

29. Badieritakis, E.; Papachristos, D.; Latinopoulos, D.; Stefopoulou, A.; Kolimenakis, A.; Bithas, K.; Patsoula, E.; Beleri, S.; Maselou, D.; Balatsos, G.; et al. Aedes Albopictus (Skuse, 1895) (Diptera: Culicidae) in Greece: 13 Years of Living with the Asian Tiger Mosquito. Parasitol Res 2018, 117, 453–460, doi:10.1007/s00436-017-5721-6.

30. Vakali, A.; Beleri, S.; Tegos, N.; Fytrou, A.; Mpimpa, A.; Sergentanis, T.N.; Pervanidou, D.; Patsoula, E. Entomological Surveillance Activities in Regions in Greece: Data on Mosquito Species Abundance and West Nile Virus Detection in Culex Pipiens Pools (2019–2020). Tropical Medicine and Infectious Disease 2023, 8, 1, doi:10.3390/tropicalmed8010001.

31. Angelini, R.; Finarelli, A.C.; Angelini, P.; Po, C.; Petropulacos, K.; Silvi, G.; Macini, P.; Fortuna, C.; Venturi, G.; Magurano, F.; et al. Chikungunya in North-Eastern Italy: A Summing up of the Outbreak. Euro Surveill 2007, 12, E071122.2, doi:10.2807/esw.12.47.03313-en.

32. Grandadam, M.; Caro, V.; Plumet, S.; Thiberge, J.-M.; Souarès, Y.; Failloux, A.-B.; Tolou, H.J.; Budelot, M.; Cosserat, D.; Leparc-Goffart, I.; et al. Chikungunya Virus, Southeastern France - Volume 17, Number 5—May 2011 - Emerging Infectious Diseases Journal - CDC., doi:10.3201/eid1705.101873.

33. Vega-Rua, A.; Zouache, K.; Caro, V.; Diancourt, L.; Delaunay, P.; Grandadam, M.; Failloux, A.-B. High Efficiency of Temperate Aedes Albopictus to Transmit Chikungunya and Dengue Viruses in the Southeast of France. PLOS ONE 2013, 8, e59716, doi:10.1371/journal.pone.0059716.

34. Delisle, E.; Rousseau, C.; Broche, B.; Leparc-Goffart, I.; L’Ambert, G.; Cochet, A.; Prat, C.; Foulongne, V.; Ferré, J.B.; Catelinois, O.; et al. Chikungunya Outbreak in Montpellier, France, September to October 2014. Eurosurveillance 2015, 20, 21108, doi:10.2807/1560-7917.ES2015.20.17.21108.

35. Calba, C.; Guerbois-Galla, M.; Franke, F.; Jeannin, C.; Auzet-Caillaud, M.; Grard, G.; Pigaglio, L.; Decoppet, A.; Weicherding, J.; Savaill, M.-C.; et al. Preliminary Report of an Autochthonous Chikungunya Outbreak in France, July to September 2017. Eurosurveillance 2017, 22, 17, doi:10.2807/1560-7917.ES.2017.22.39.17-00647.

36. Calba, C.; Guerbois-Galla, M.; Franke, F.; Jeannin, C.; Auzet-Caillaud, M.; Grard, G.; Pigaglio, L.; Cadiou, B.; de Lamballerie, X.; Paty, M.-C.; et al. Investigation of an Autochthonous Chikungunya Outbreak, July–September 2017, France. Revue d’Épidémiologie et de Santé Publique 2018, 66, S387–S388, doi:10.1016/j.respe.2018.05.410.

37. Riccardo, F.; Venturi, G.; Luca, M.D.; Manso, M.D.; Severini, F.; Andrianou, X.; Fortuna, C.; Remoli, M.E.; Benedetti, E.; Caporali, M.G.; et al. Secondary Autochthonous Outbreak of Chikungunya, Southern Italy, 2017 - Volume 25, Number 11—November 2019 - Emerging Infectious Diseases Journal - CDC., doi:10.3201/eid2511.180949.

38. Venturi, G.; Luca, M.D.; Fortuna, C.; Remoli, M.E.; Riccardo, F.; Severini, F.; Toma, L.; Manso, M.D.; Benedetti, E.; Caporali, M.G.; et al. Detection of a Chikungunya Outbreak in Central Italy, August to September 2017. Eurosurveillance 2017, 22, 17, doi:10.2807/1560-7917.ES.2017.22.39.17-00646.

39. Gjenero-Margan, I.; Aleraj, B.; Krajcar, D.; Lesnikar, V.; Klobučar, A.; Pem-Novosel, I.; Kurečić-Filipović, S.; Komparak, S.; Martić, R.; Đuričić, S.; et al. Autochthonous Dengue Fever in Croatia, August–September 2010. Eurosurveillance 2011, 16, 19805, doi:10.2807/ese.16.09.19805-en.

40. Ruche, G.L.; Souarès, Y.; Armengaud, A.; Peloux-Petiot, F.; Delaunay, P.; Desprès, P.; Lenglet, A.; Jourdain, F.; Leparc-Goffart, I.; Charlet, F.; et al. First Two Autochthonous Dengue Virus Infections in Metropolitan France, September 2010. Eurosurveillance 2010, 15, 19676, doi:10.2807/ese.15.39.19676-en.

41. Gossner, C.M.; Fournet, N.; Frank, C.; Fernández-Martínez, B.; Manso, M.D.; Dias, J.G.; Valk, H. de Dengue Virus Infections among European Travellers, 2015 to 2019. Eurosurveillance 2022, 27, 2001937, doi:10.2807/1560-7917.ES.2022.27.2.2001937.

42. Fournet, N.; Voiry, N.; Rozenberg, J.; Bassi, C.; Cassonnet, C.; Karch, A.; Durand, G.; Grard, G.; Modenesi, G.; Lakoussan, S.-B.; et al. A Cluster of Autochthonous Dengue Transmission in the Paris Region – Detection, Epidemiology and Control Measures, France, October 2023. Eurosurveillance 2023, 28, 2300641, doi:10.2807/1560-7917.ES.2023.28.49.2300641.

43. ECDC Local Transmission of Dengue Virus in Mainland EU/EEA, 2010-Present Available online: https://www.ecdc.europa.eu/en/all-topics-z/dengue/surveillance-and-disease-data/autochthonous-transmission-dengue-virus-eueea (accessed on 18 December 2024).

44. Carli, G.D.; Carletti, F.; Spaziante, M.; Gruber, C.E.M.; Rueca, M.; Spezia, P.G.; Vantaggio, V.; Barca, A.; Liberato, C.D.; Romiti, F.; et al. Outbreaks of Autochthonous Dengue in Lazio Region, Italy, August to September 2023: Preliminary Investigation. Eurosurveillance 2023, 28, 2300552, doi:10.2807/1560-7917.ES.2023.28.44.2300552.

45. Sacco, C.; Liverani, A.; Venturi, G.; Gavaudan, S.; Riccardo, F.; Salvoni, G.; Fortuna, C.; Marinelli, K.; Marsili, G.; Pesaresi, A.; et al. Autochthonous Dengue Outbreak in Marche Region, Central Italy, August to October 2024. Eurosurveillance 2024, 29, 2400713, doi:10.2807/1560-7917.ES.2024.29.47.2400713.

46. Kumar, N.P.; Rajavel, A.R.; Natarajan, R.; Jambulingam, P. DNA Barcodes Can Distinguish Species of Indian Mosquitoes (Diptera: Culicidae). J Med Entomol 2007, 44, 1–7, doi:10.1603/0022-2585(2007)44[1:dbcdso]2.0.co;2.

47. Cywinska, A.; Hunter, F.F.; Hebert, P.D.N. Identifying Canadian Mosquito Species through DNA Barcodes. Med Vet Entomol 2006, 20, 413–424, doi:10.1111/j.1365-2915.2006.00653.x.

48. Hemmerter, S.; Slapeta, J.; Beebe, N.W. Resolving Genetic Diversity in Australasian Culex Mosquitoes: Incongruence between the Mitochondrial Cytochrome c Oxidase I and Nuclear Acetylcholine Esterase 2. Mol Phylogenet Evol 2009, 50, 317–325, doi:10.1016/j.ympev.2008.11.016.

49. Wang, G.; Li, C.; Guo, X.; Xing, D.; Dong, Y.; Wang, Z.; Zhang, Y.; Liu, M.; Zheng, Z.; Zhang, H.; et al. Identifying the Main Mosquito Species in China Based on DNA Barcoding. PLOS ONE 2012, 7, e47051, doi:10.1371/journal.pone.0047051.

50. Pradeep Kumar, N.; Krishnamoorthy, N.; Sahu, S.S.; Rajavel, A.R.; Sabesan, S.; Jambulingam, P. DNA Barcodes Indicate Members of the Anopheles Fluviatilis (Diptera: Culicidae) Species Complex to Be Conspecific in India. Mol Ecol Resour 2013, 13, 354–361, doi:10.1111/1755-0998.12076.

51. Patsoula, E.; Samanidou-Voyadjoglou, A.; Spanakos, G.; Kremastinou, J.; Nasioulas, G.; Vakalis, N.C. Molecular and Morphological Characterization of Aedes Albopictus in Northwestern Greece and Differentiation from Aedes Cretinus and Aedes Aegypti. J Med Entomol 2006, 43, 40–54, doi:10.1093/jmedent/43.1.40.

52. Zheng, M.-L.; Zhang, D.-J.; Damiens, D.D.; Lees, R.S.; Gilles, J.R.L. Standard Operating Procedures for Standardized Mass Rearing of the Dengue and Chikungunya Vectors Aedes Aegypti and Aedes Albopictus (Diptera: Culicidae) - II - Egg Storage and Hatching. Parasites & Vectors 2015, 8, 348, doi:10.1186/s13071-015-0951-x.

53. Darsie, R.F.; Samanidou-Voyadjoglou, A. Keys for the Identification of the Mosquitoes of Greece. J Am Mosq Control Assoc 1997, 13, 247–254.

54. Becker, N.; Petric, D.; Zgomba, M.; Boase, C.; Madon, M.; Dahl, C.; Kaiser, A. Mosquitoes and Their Control; Springer: Berlin, Heidelberg, 2010; ISBN 978-3-540-92873-7.

55. Patsoula, E.; Beleri, S.; Vakali, A.; Pervanidou, D.; Tegos, N.; Nearchou, A.; Daskalakis, D.; Mourelatos, S.; Hadjichristodoulou, C. Records of Aedes Albopictus (Skuse, 1894) (Diptera; Culicidae) and Culex Tritaeniorhynchus (Diptera; Culicidae) Expansion in Areas in Mainland Greece and Islands. Vector-Borne and Zoonotic Diseases 2017, 17, 217–223, doi:10.1089/vbz.2016.1974.

56. Simon, C.; Frati, F.; Beckenbach, A.; Crespi, B.; Liu, H.; Flook, P. Evolution, Weighting, and Phylogenetic Utility of Mitochondrial Gene Sequences and a Compilation of Conserved Polymerase Chain Reaction Primers. Annals of the Entomological Society of America 1994, 87, 651–701, doi:10.1093/aesa/87.6.651.

57. Molecular Genetic Analysis of Populations: A Practical Approach; Hoelzel, A.R., Ed.; The practical approach series; 2nd ed.; IRL Press at Oxford University Press: Oxford, 1998; ISBN 978-0-19-963634-1.

58. Milligan, B. g. Total DNA Isolation. In Molecular Genetic Analysis of Populations: A Practical Approach; Hoelzel, A.R., Ed.; Oxford University Press, 1998; p. 0 ISBN 978-0-19-963634-1.

59. Folmer, O.; Black, M.; Hoeh, W.; Lutz, R.; Vrijenhoek, R. DNA Primers for Amplification of Mitochondrial Cytochrome c Oxidase Subunit I from Diverse Metazoan Invertebrates. Mol Mar Biol Biotechnol 1994, 3, 294–299.

60. Beebe, N.W.; Ambrose, L.; Hill, L.A.; Davis, J.B.; Hapgood, G.; Cooper, R.D.; Russell, R.C.; Ritchie, S.A.; Reimer, L.J.; Lobo, N.F.; et al. Tracing the Tiger: Population Genetics Provides Valuable Insights into the Aedes (Stegomyia) Albopictus Invasion of the Australasian Region. PLoS Negl Trop Dis 2013, 7, e2361, doi:10.1371/journal.pntd.0002361.

61. Delatte, H.; Toty, C.; Boyer, S.; Bouetard, A.; Bastien, F.; Fontenille, D. Evidence of Habitat Structuring Aedes Albopictus Populations in Réunion Island. PLOS Neglected Tropical Diseases 2013, 7, e2111, doi:10.1371/journal.pntd.0002111.

62. Ashfaq, M.; Hebert, P.D.N.; Mirza, J.H.; Khan, A.M.; Zafar, Y.; Mirza, M.S. Analyzing Mosquito (Diptera: Culicidae) Diversity in Pakistan by DNA Barcoding. PLOS ONE 2014, 9, e97268, doi:10.1371/journal.pone.0097268.

63. Medley, K.A.; Jenkins, D.G.; Hoffman, E.A. Human-Aided and Natural Dispersal Drive Gene Flow across the Range of an Invasive Mosquito. Mol Ecol 2015, 24, 284–295, doi:10.1111/mec.12925.

64. Manni, M.; Gomulski, L.M.; Aketarawong, N.; Tait, G.; Scolari, F.; Somboon, P.; Guglielmino, C.R.; Malacrida, A.R.; Gasperi, G. Molecular Markers for Analyses of Intraspecific Genetic Diversity in the Asian Tiger Mosquito, Aedes Albopictus. Parasites & Vectors 2015, 8, 188, doi:10.1186/s13071-015-0794-5.

65. Brown, W.M.; George, M.; Wilson, A.C. Rapid Evolution of Animal Mitochondrial DNA. Proc Natl Acad Sci U S A 1979, 76, 1967–1971, doi:10.1073/pnas.76.4.1967.

66. Tantely, M.L.; Tortosa, P.; Alout, H.; Berticat, C.; Berthomieu, A.; Rutee, A.; Dehecq, J.-S.; Makoundou, P.; Labbé, P.; Pasteur, N.; et al. Insecticide Resistance in Culex Pipiens Quinquefasciatus and Aedes Albopictus Mosquitoes from La Réunion Island. Insect Biochem Mol Biol 2010, 40, 317–324, doi:10.1016/j.ibmb.2010.02.005.

67. Kawada, H.; Maekawa, Y.; Abe, M.; Ohashi, K.; Ohba, S.; Takagi, M. Spatial Distribution and Pyrethroid Susceptibility of Mosquito Larvae Collected from Catch Basins in Parks in Nagasaki City, Nagasaki, Japan. Jpn J Infect Dis 2010, 63, 19–24.

68. Gasperi, G.; Bellini, R.; Malacrida, A.R.; Crisanti, A.; Dottori, M.; Aksoy, S. A New Threat Looming over the Mediterranean Basin: Emergence of Viral Diseases Transmitted by Aedes Albopictus Mosquitoes. PLoS Negl Trop Dis 2012, 6, e1836, doi:10.1371/journal.pntd.0001836.

69. Bonizzoni, M.; Gasperi, G.; Chen, X.; James, A.A. The Invasive Mosquito Species Aedes Albopictus: Current Knowledge and Future Perspectives. Trends Parasitol 2013, 29, 460–468, doi:10.1016/j.pt.2013.07.003.

